# SARM1 promotes axonal, synaptic, and mitochondrial pathologies in Charcot-Marie-Tooth disease type 2A

**DOI:** 10.1101/2022.05.17.492364

**Authors:** Yurie Sato-Yamada, Amy Strickland, Yo Sasaki, Joseph Bloom, Aaron DiAntonio, Jeffrey Milbrandt

## Abstract

Charcot-Marie-Tooth disease (CMT) type 2A is an axonal neuropathy caused by mutations in the mitofusin 2 (*MFN2*) gene. *MFN2* mutations result in profound mitochondrial abnormalities, but the mechanism underlying axonal pathology is unknown. SARM1, the central executioner of axon degeneration, can induce neuropathy and is activated by dysfunctional mitochondria. We tested the role of SARM1 in a rat model carrying a dominant CMT2A mutation (*Mfn2^H361Y^*) that exhibits progressive dying-back axonal degeneration, NMJ abnormalities, muscle atrophy, and mitochondrial abnormalities, all hallmarks of the human disease. We generated *Sarm1* knockout and *Mfn2^H361Y^, Sarm1* double mutant rats and find that deletion of SARM1 rescues axonal, synaptic, and muscle phenotypes, demonstrating that SARM1 induces much of the neuropathology in this model. Despite the presence of mutant Mfn2 protein in these double mutant rats, loss of SARM1 also dramatically suppressed many mitochondrial defects, including the number, size, and cristae density defects of synaptic mitochondria. This surprising finding indicates that dysfunctional mitochondria activate SARM1, and activated SARM1 feeds back on mitochondria to exacerbate mitochondrial pathology. As such, this work identifies SARM1 inhibition as an exciting therapeutic candidate for the treatment of CMT2A and other neurodegenerative diseases with prominent mitochondrial pathology.

## Introduction

Charcot-Marie-Tooth disease 2A (CMT2A) is a common hereditary motor and sensory neuropathy of the peripheral nervous system that is characterized by progressive, length-dependent axonal degeneration without myelin involvement that predominantly affects the distal limbs. CMT2A tends to have an earlier onset and faster progression than most CMTs, leaving many patients non-ambulatory as children (1, 2). As with all CMTs, there are no disease-modifying treatments. CMT2A is caused by mutations in the Mytofusin2 (*MFN2*) gene. MFN2 is a nuclear-encoded dynamin-like GTPase residing in the outer membrane of mitochondria that plays a critical role in mitochondrial fusion, but also promotes mitochondrial mobility, mitophagy, and inter-organelle calcium signaling (3). In CMT2A patients, neuronal mitochondria have morphological and functional abnormalities (2, 4, 5), but how these mitochondrial defects result in dying-back axon loss is unknown.

SARM1 is the central executioner of the programmed axon destruction pathway (6, 7), and so is a candidate to mediate axon loss in CMT2A. SARM1 is an enzyme that, when activated, cleaves the essential metabolic co-factor NAD^+^ (8), inducing a stereotyped local metabolic collapse with loss of ATP, defects in mitochondrial motility and depolarization, followed by calcium influx and ultimately axon fragmentation (9). Loss of SARM1 blocks axon degeneration in mouse models of axotomy (10, 11), traumatic brain injury (12), and glaucoma (13). Notably, loss of SARM1 also preserves axons in multiple models of chemotherapeutic and metabolic peripheral neuropathy (14–18). Moreover, in a recently described mouse model of a human motor neuropathy caused by loss of the SARM1 inhibitor NMNAT2, SARM1 mediates a slowly-progressive motor-predominant neuropathy with axon loss and muscle atrophy (19), both hallmarks of CMT2A. Finally, genetic variants of *SARM1* that encode constitutively-active enzymes are enriched in patients with ALS and other motor neuropathies (20, 21). In addition to the strong evidence that SARM1 induces axon loss in neuropathies, there is also a wealth of data demonstrating that mitochondrial dysfunction activates SARM1 (22–25). Thus, we hypothesized that SARM1 may be activated in CMT2A neurons and induce axon loss. If SARM1 promotes neuropathology in CMT2A, then this would open new possibilities for treatment as both chemical and gene therapy SARM1inhibitors block axon loss (26–28).

The role of SARM1 in neurodegenerative disease has primarily been tested in the mouse. While mouse models of CMT2A exist, they have significant limitations (3). Overexpression of human CMT2A-associated *MFN2* variants using different transgenic systems gives significant phenotypic heterogeneity (29–33), while knock-in of a pathogenic mutation into the endogenous mouse *Mfn2* locus does not cause axonal defects (34). To overcome these obstacles, here we analyze the recently described Mfn2 knock-in rat model (35) carrying the strong pathogenic human mutation H361Y (36). This model recapitulates many aspects of human CMT2A (37). We analyzed the neuropathology in this rat model and find a progressive motor axonopathy with neuromuscular junction (NMJ) defects and muscle atrophy. In addition, we observe numerous mitochondrial defects including fewer mitochondria at synapses, abnormal accumulation of aggregated mitochondria near nodes of Ranvier, and defects of mitochondrial shape and cristae density most prominent in distal portions of the nerve. We generated a *Sarm1* knockout rat as well as the *Mfn2^H361Y^,Sarm1* double mutant rat in order to test the role of SARM1 in CMT2A. We demonstrate that SARM1 is required for axon degeneration, NMJ defects, and hind limb muscle atrophy in the *Mfn2^H361Y^* rat. Therefore, SARM1 is a key driver of neuropathology in this model of CMT2A. Since mutant Mfn2 protein is present in the *Mfn2^H361Y^,Sarm1* rat, we expected that mitochondrial defects would persist. Instead, SARM1 deletion also suppresses defects in mitochondrial localization, size, number, and cristae density. In addition, deletion of SARM1 improves mitochondrial axon transport in an *in vitro* assay. These surprising findings demonstrate that dysfunctional mitochondria activate SARM1 to induce the major neuropathological defects in this CMT2A model, and that activated SARM1 feeds back onto mitochondria, exacerbating their dysfunction. Hence, SARM1 inhibition is a compelling therapeutic candidate for the treatment of CMT2A and potentially the many other neurodegenerative diseases characterized by mitochondrial dysfunction.

## Results

### *Mfn2^H361Y/+^* rats recapitulate CMT2A neuromuscular phenotypes

CMT2A patients exhibit loss of vibratory sense as well as motor weakness that is predominantly in the distal lower limb and rapidly progresses throughout the first decade of life (1). Sural nerve pathology demonstrates loss of large myelinated fibers with regenerating clusters, but no myelin abnormalities (5). Mitochondrial abnormalities include swelling and dissolution of cristae, as well as aggregation within the axon (4, 37–39). To model this disease, the *Mfn2^H361Y^* mutation, which causes severe early onset disease in patients, was introduced into the rat genome using CRISPR/Cas9. This MFN2A model rat develops progressive functional abnormalities and loss of myelinated axons (35). To investigate the neuronal pathologies in *Mfn2^H361Y/+^* rats, we examined the hind limb nerves and muscles of *Mfn2^H361Y/+^* rats at 6 and 12 months of age. Axons in the sciatic nerve were intact at 6 months of age but by 12 months there was severe axon loss in the distal sciatic nerve (Fig. 1A, 2C). Morphometric analysis of axon diameters in 12-month-old *Mfn2^H361Y/+^* rats revealed a decrease in large caliber axons that was mirrored by an increase in the proportion of small fibers (Fig. 1B). This axon size distribution is consistent with a motor neuropathy (40), and is similar to that observed in CMT2A human nerve (37). CMT2A patients exhibit symptoms mainly in the lower leg, attributed to more extensive distal axonal degeneration. Tellingly, no axonal defects in proximal sciatic nerves were observed in *Mfn2^H361Y/+^rats* (Fig. 1A), consistent with a dying back neuropathy. Another pathological signature in CMT2A peripheral nerves is presence of clusters of regenerating axons along with atrophied axons and onion bulb structures, cardinal evidence of repetitive axonal degeneration and regeneration (37). To detect regenerating axons in distal hind limb nerves, we performed immunostaining for STMN2, a marker of regenerating axons (41). Many STMN2-positive axons were detected in 12-month-old *Mfn2^H361Y/+^* tibial nerves, while only a few STMN2-labeled axons were seen in the nerves of 6-month-old *Mfn2^H361Y/+^* or WT rats (Fig. 1C).

**Figure 1.**
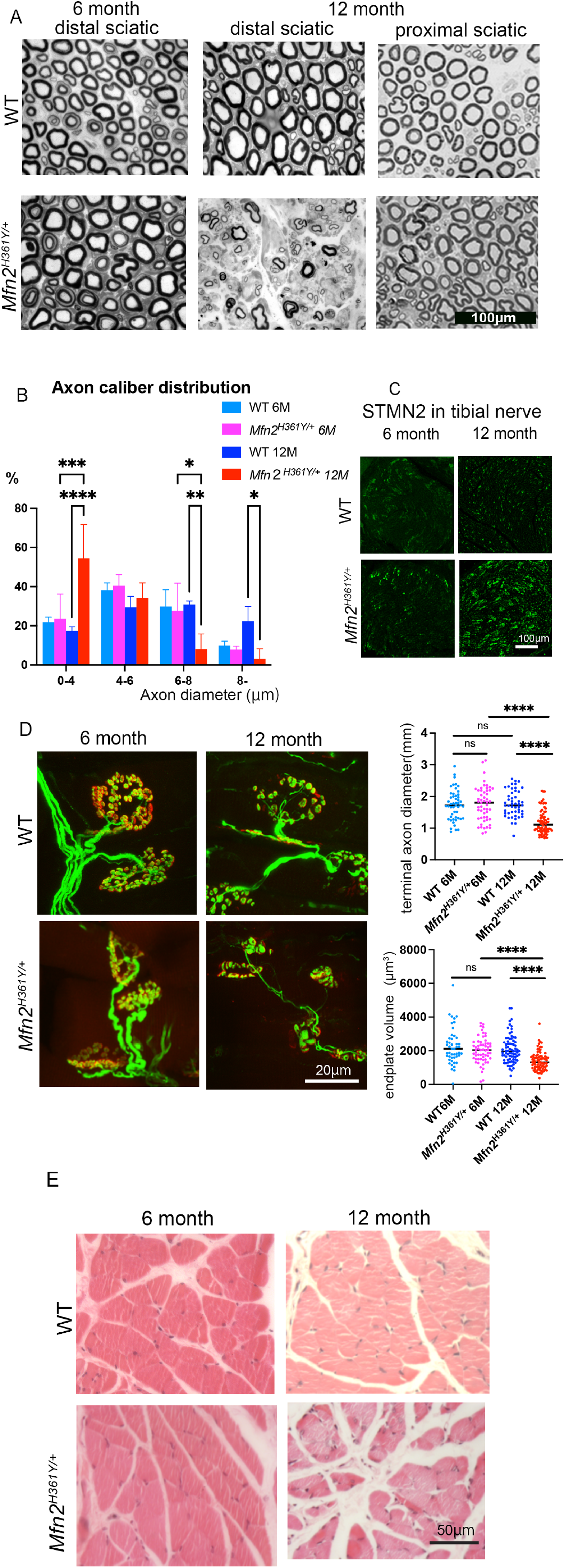
*Mfn2* H361Y mutation causes progressive neurodegeneration and muscle wasting. (A) Toluidine blue-stained cross sections of sciatic nerves from WT and *Mfn2^H361Y/+^* rats at 6 and 12 months of age. (B) Distribution of axonal diameters of distal sciatic nerves in WT and *Mfn2^H361Y/+^* rats at 6 and 12 months of age (n = 3). *p<0.05, **p<0.01, ***p<0.005, ****p<0.001(C) STMN2 in tibial nerves of WT and *Mfn2^H361Y/+^* rats at 6 and 12 months of age. (D) Representative images of NMJs in WT and *Mfn2 ^H361Y/+^* rats at 6 and 12 months of age stained in green to detect the synaptic vesicle marker SV2A (synaptic vesicle glycoprotein 2A) and axon marker NEFM (neurofilament medium chain) and in red with the post-synaptic endplate marker bungarotoxin. The upper graph exhibits terminal axon diameters and the lower graph exhibits endplate volumes in WT and *Mfn2 ^H361Y/+^* rats at 6 and 12 months of age (n = 3-4). ****p< 0.001. (E) Cross sections of lumbrical muscles stained with hematoxylin and eosin from WT and *Mfn2 ^H361Y/+^* rats at 6 and 12 months of age.

**Figure 2.**
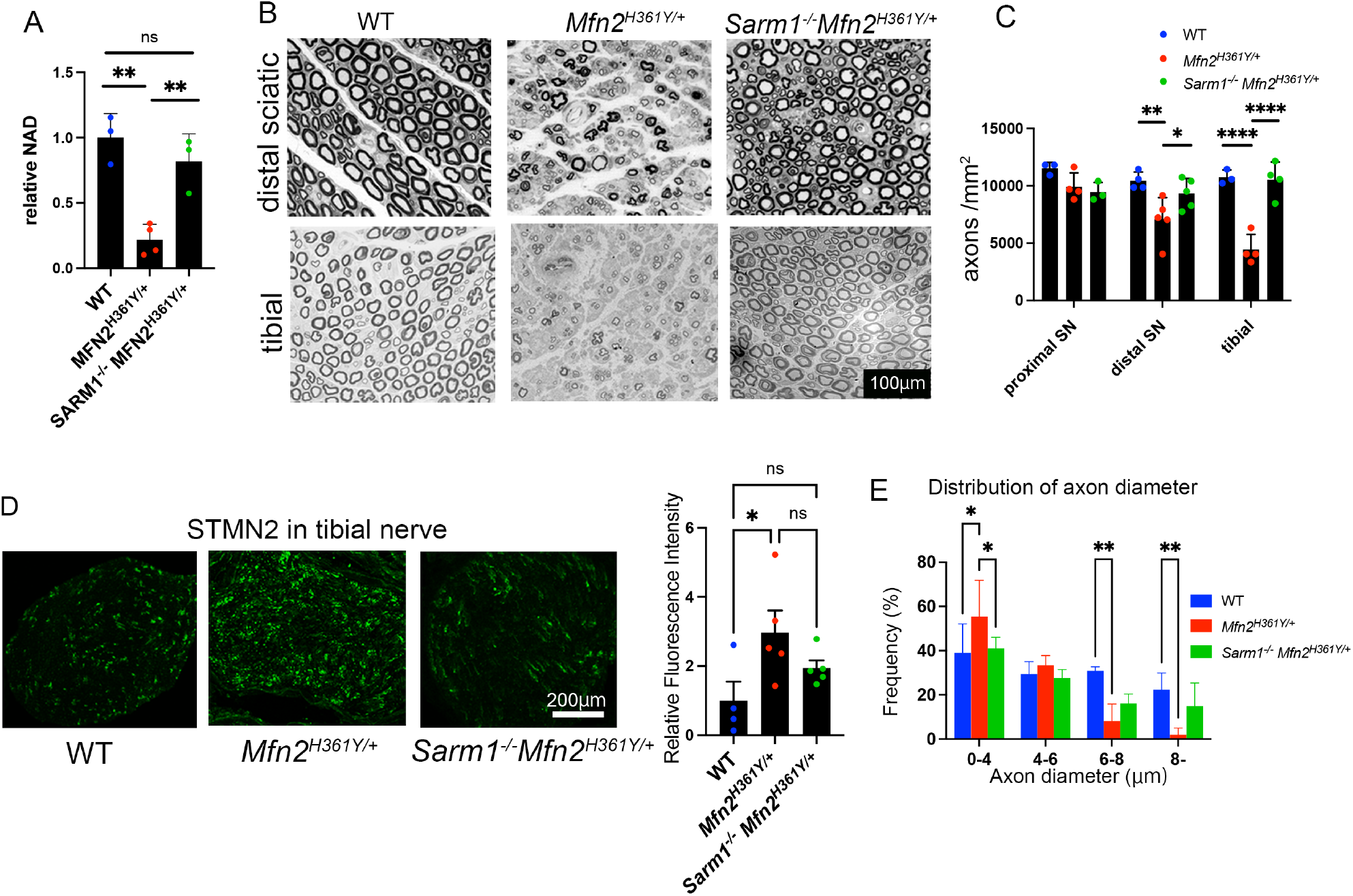
*Sarm1* deletion protects nerves from axonal degeneration in *Mfn2^H361Y/+^* rats. (A) Relative NAD^+^ level in tibial nerves of 12 month old WT, *Mfn2^H361Y/+^*, and *Sarm1^-/-^Mfn2^H361Y/+^* rats. Values normalized to WT (n = 3-4). **p<0.01. (B) Toluidine blue-stained cross sections of distal sciatic nerve and tibial nerve from WT, *Mfn2 ^H361Y/+^*, and *Sarm1^-/-^Mfn2^H361Y/+^* rats. (C) The number of axons per mm^2^ in proximal sciatic nerve, distal sciatic nerve, and tibial nerve from WT, *Mfn2^H361Y/+^*, and *Sarm1^-/-^Mfn2^H361Y/+^* rats (n=3-4). *p<0.05, **p<0.01, ****p<0.001. (D) STMN2 in tibial nerve from WT, *Mfn2^H361Y/+^*, and *Sarm1^-/-^Mfn2^H361Y/+^* rats. Graph shows the relative fluorescence intensity of images for each genotype (n = 3). *p<0.05. (E) Distribution of axonal diameters of distal sciatic nerve from WT, *Mfn2 ^H361Y/+^*, and *Sarm1^-/-^Mfn2^H361Y/+^* rats (n = 3). *p<0.05, **p<0.01

A number of motor-predominant neurodegenerative diseases exhibit abnormalities in the structure of the neuromuscular junction (NMJ), an early and important site of neuropathology in type 2 axonal CMT (42). We stained NMJs in lumbrical muscles with antibodies against neurofilament (axon marker) and SV2 (synapse vesicle marker), and α-bungarotoxin (post-synaptic marker). Morphological abnormalities found in the NMJs of 12-month *Mfn2^H361Y/+^* rats included thin terminal axons and shrunken endplates (Fig. 1D). Of note, NMJs innervated by abnormally thin terminal axons have been associated with models of motor neuron disorders including amyotrophic lateral sclerosis (43) and spinal and bulbar muscular atrophy (44). Lower leg muscle atrophy is a typical clinical symptom of CMT2A (37), so we searched for pathology in lumbrical muscles of hind paws of 12-month old *Mfn2^H361Y/+^* rats. We found smaller muscle fascicles (WT=435.7 *μ* m^2^ vs. *Mfn2^H361Y/+^*=356.2 μm^2^, n=3, p=0.045, Fig. 1E) in the mutant rats, indicating the *Mfn2^H361Y^* mutation causes progressive neuropathy in the hind limb. The axonal degeneration, NMJ pathology and muscle atrophy in these *Mfn2^H361Y/+^* rats suggest they are a faithful model of human CMT2A.

### *Sarm1* knockout prevents *Mfn2^H361Y/+^*-associated axon and muscle defects

SARM1 is an essential component of the programmed axon destruction pathway. Activation of SARM1 triggers axonal degeneration, and the deletion of *SARM1* protects axons from acute injury-induced Wallerian degeneration and toxic and metabolic peripheral neuropathy (10, 11, 14–16, 45); however, whether SARM1 is involved in progressive neurodegenerative conditions like CMT2A is untested.

To investigate whether SARM1 contributes to axonal degeneration in CMT2A, we generated *Sarm1* mutant rats using CRISPR gene editing (Supplementary Fig. 1A). The deletion in *Sarm1* and consequent loss of SARM1 protein were confirmed by DNA sequencing and western blotting of brain lysates. *Sarm1 KO* rats were further subjected to a sciatic nerve transection assay to test the axon protective phenotype of this *Sarm1* loss-of-function allele. Plastic sections of distal sciatic nerve from *Sarm1* KO and wildtype rats were analyzed 7 days-post transection (Supplemental Fig.1C, and showed robust protection of the transected axons as has been previously reported in *Sarm1* KO mice (10, 11).

The *Sarm1* KO rats were then crossed with *Mfn2^H361Y/+^* rats to test whether loss of SARM1 prevented the pathology observed in this CMT2A model. To investigate whether SARM1 is activated in the nerves of *Mfn2^H361Y/+^* rats, we first examined levels of NAD^+^, the SARM1 substrate (8). When SARM1 is activated, it cleaves NAD^+^, and NAD^+^ levels decrease. LC/MS-MS revealed a decline of NAD^+^ in *Mfn2^H361Y/+^* rat tibial nerves, whereas levels remained equivalent to WT in *Sarm1^-/-^Mfn2^H361Y/+^* double mutant nerves (Fig. 2A). Crucially, the dramatic axonal loss and increased number of regenerating axons (marked by STMN2 staining) observed in *Mfn2^H361Y/+^* nerves are completely abrogated in 12-month-old *Sarm1^-/-^Mfn2^H361Y/+^* rats (Fig. 2B-D). The altered distribution of axonal diameters in *Mfn2^H361Y/+^* nerves was also suppressed in *Sarm1^-/-^Mfn2^H361Y/+^* rats (Fig. 2E), indicating the preservation of large caliber axons with deletion of *Sarm1*.

Along with the axonal deficits, the distal muscles in *Mfn2^H361Y/+^* rats show severe atrophy (Fig 1E). We further investigated these muscle defects and tested whether SARM1 deletion also improves the loss of muscle integrity caused by the *Mfn2^H361Y/+^* mutation. We found atrophy of isolated myofibers, typically seen after loss of innervation (46), in the tibialis anterior (TA) muscles (Fig. 3A). Muscle fascicles with centered, abnormal localization of nuclei (Fig. 3A arrowhead) were also detected, indicating a history of repeated muscle fiber atrophy and regeneration in *Mfn2^H361Y/+^* rats (47). Other muscle pathologies associated with motor neuropathy (48) and denervated muscle fibers include changes in muscle fiber diameter (49). We used laminin immunohistochemistry to highlight the muscle fibers and analyzed their cross-sectional area (CSA). We found an increased number of small caliber muscle fibers (Fig. 3B arrows) and a corresponding decrease of large caliber muscle fibers in *Mfn2^H361Y/+^* rats (Fig. 3C). Muscle denervation is often associated with increased fibrosis due to excessive accumulation of extracellular matrix that replaces functional tissue (50). Sirius red staining to detect collagen fibers showed increased collagen fiber content in *Mfn2^H361Y/+^* muscles, with more severe fibrosis and atrophy in lumbrical muscles than in TA muscles, suggesting that the denervation and associated muscle atrophy are more severe at distal than proximal nerve endings (Fig. 3D). In contrast to these results, muscles from *Sarm1^-/-^Mfn2^H361Y/+^* rats showed no change in muscle fiber caliber or evidence of fibrosis (Fig. 3A-D). Together, these results are consistent with a distal-predominant nerve and muscle pathologies in this CMT2A model. Further, they indicate that the axon degeneration due to mitochondrial dysfunction caused by *Mfn2* mutation requires SARM1 activity, demonstrating that SARM1 plays a crucial role in chronic, progressive neuropathy in addition to its known roles in acute injury.

**Figure 3.**
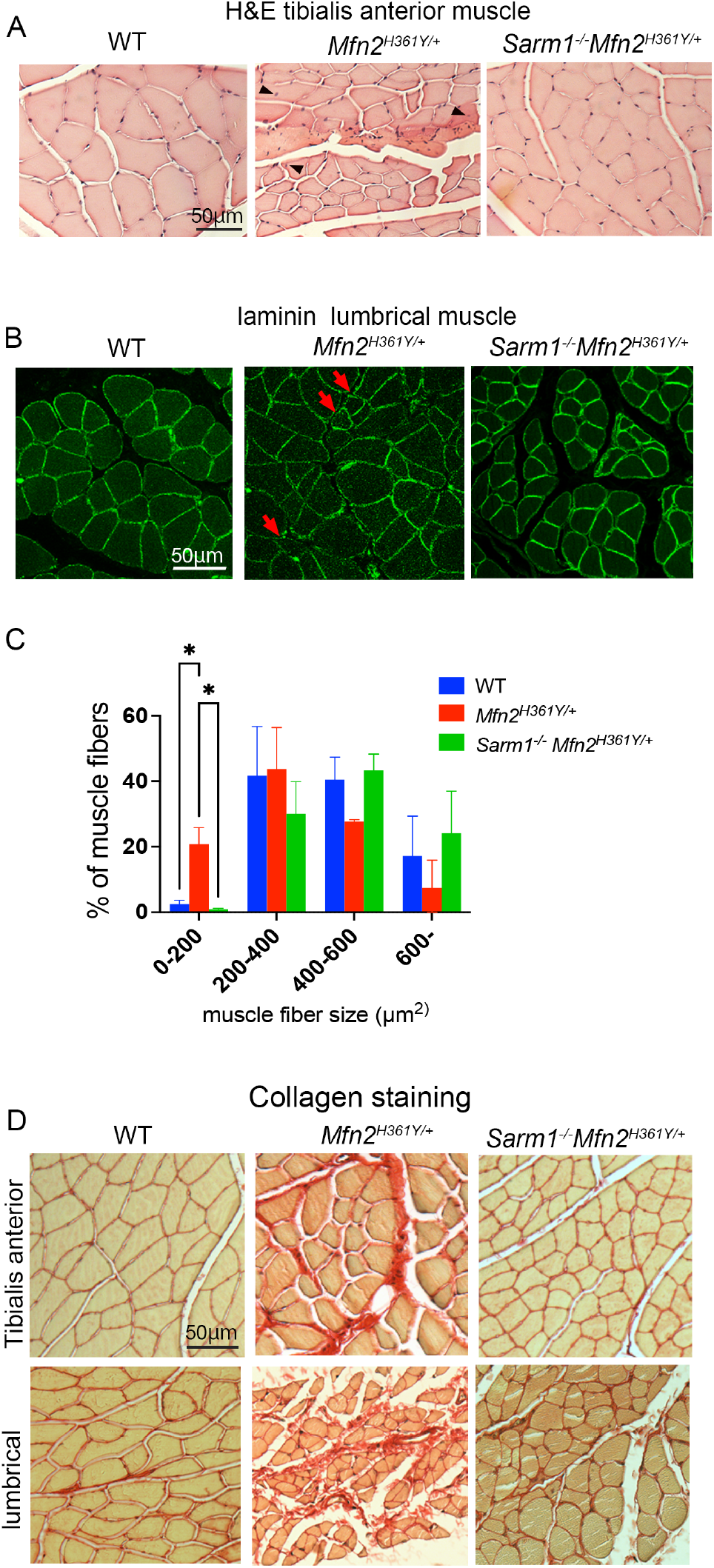
*Sarm1* deletion prevents muscle atrophy in *Mfn2 ^H361Y/+^* rats. (A) Cross sections of tibial anterior muscle stained with hematoxylin and eosin from WT, *Mfn2 ^H361Y/+^*, and *Sarm1^-/-^Mfn2^H361Y/+^* rats. Arrowheads indicate nuclei positioned in the center of myofibers, a feature of regenerated muscle. (B) Cross sections of tibial anterior muscle immunostained for laminin from WT, *Mfn2^H361Y+^*, and *Sarm1^-/-^Mfn2^H361Y/+^* rats. Arrows indicate small atrophied muscle fascicles. (C) Distribution of cross-sectional area of tibial anterior muscle fascicles from WT, *Mfn2^H361Y/+^*, and *Sarm1^-/-^Mfn2^H361Y/+^* rats (n = 3). *p < 0.05. (D) Cross sections of tibial anterior and lumbrical muscle stained with picrosirius red from WT, *Mfn2^H361Y/+^*, and *Sarm1^-/-^Mfn2^H361Y/+^* rats.

### *Mfn2^H361Y/+^* NMJ abnormalities are dependent on SARM1 activity

NMJ morphological abnormalities leading to loss of integrity vary among motor neuron diseases and are correlated with symptom severity (42, 51, 52). To examine the influence of SARM1 deletion on mutant MFN2-derived alterations in NMJ morphology, we first classified lumbrical muscle NMJs into three categories defined previously by others (53, 54) (Fig. 4A): 1) ‘Normal NMJs’ with typical axon diameters at terminals (>1.8 μm) and synaptic terminal branches closely opposed to acetylcholine receptor-rich endplates; 2) ‘Thin NMJs’ with narrow axonal diameters (<1.8 μm) and a mixture of retained (arrow) or eliminated (arrowhead) junctions; and 3) ‘Denervated NMJs’ with α-BTX labeled postsynaptic sites lacking pre-synaptic structures or terminal axons. *Mfn2^H361Y/+^* rats exhibited few normal NMJs (*Mfn2^H361Y/+^* 20.8%, WT 71.6%), with most of their NMJs classified as thin (70.5%) and a few as denervated (8.8%).

**Figure 4.**
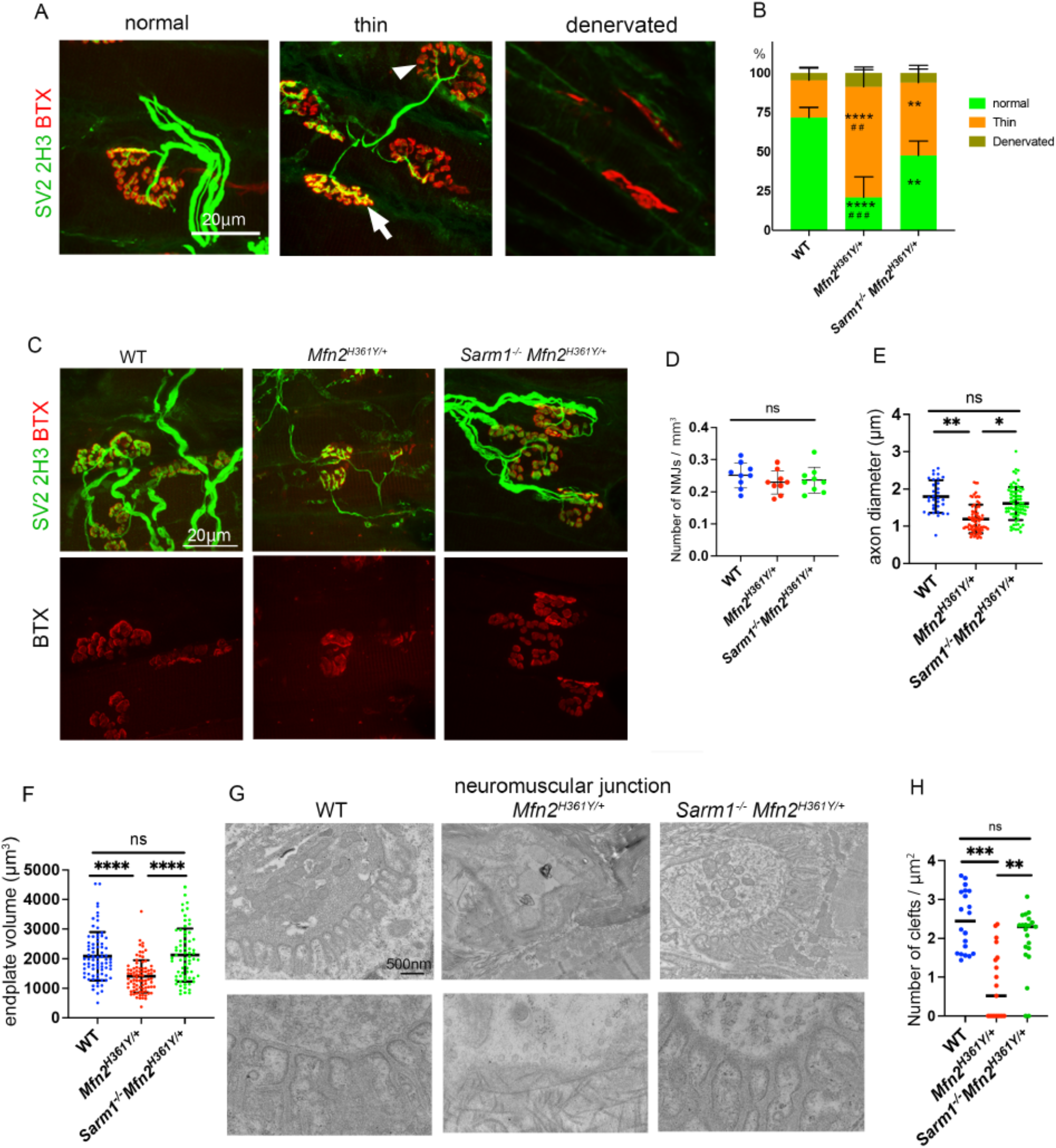
*Sarm1* deletion protects NMJs from degenerating in *Mfn2^H361Y/+^* rats. (A) Representative images of *Mfn2 ^H361Y/+^* NMJs with various morphologies stained to detect the synaptic vesicle marker SV2A (SV2), the axon marker NEFM (2H3), and post-synaptic endplate marker α-bungarotoxin (BTX). Arrow indicates an endplate innervated by a thin axon and arrowhead indicates an endplate which lacks a pre-synaptic structure. (B) Percentage of each NMJ category in WT, *Mfn2^H361Y/+^*, and *Sarm1^-/-^Mfn2^H361Y/+^* lumbrical muscles (n = 4). * indicates statistical significance comparing WT to *Mfn2^H361Y/+^* or *Sarm1^-/-^Mfn2^H361Y+^*, # indicates statistical significance comparing *Mfn2^H361Y/+^* to *Sarm1^-/-^Mfn2^H361Y/+^*. ** ## p<0.01, ### p<0.005, ****p<0.001, (C) Representative images of NMJs in WT, *Mfn2^H361Y/+^*, and *Sarm1^-/-^Mfn2^H361Y/+^* lumbrical muscles. (D) NMJs per mm^3^ in WT, *Mfn2^H361Y/+^*, and *Sarm1^-/-^Mfn2^H361Y/+^* lumbrical muscles (n =3-4). (E) Terminal axon diameters in WT, *Mfn2^H361Y/+^*, and *Sarm1^-/-^Mfn2^H361Y/+^* lumbrical muscles (n = 3-4). *p<0.05, **p<0.01. (F) Endplate volume in WT, *Mfn2^H361Y/+^*, and *Sarm1^-/-^Mfn2^H361Y/+^* lumbrical muscles NMJs (n = 3-4). ****p<0.001. Some data in E & F are reiterated from Fig. 1D for comparison. (G) Electron microscope images of NMJs in WT, *Mfn2^H361Y/+^*, and *Sarm1^-/-^Mfn2^H361Y+^* lumbrical muscles. (H) Synaptic clefts per μm^2^ in WT, *Mfn2^H361Y/+^*, and *Sarm1^-/-^Mfn2^H361Y/+^* lumbrical muscle NMJs (n = 3-4). **p<0.01, ***p<0.005.

In performing similar analyses of *Sarm1^-/-^Mfn2^H361Y/+^* muscles, we found that the endplates and terminal axons looked morphologically normal, with a significant decrease in the percentage of abnormal NMJs compared to *Mfn2^H361Y/+^* (thin NMJs 46.1%, denervated NMJs 6.3%) (Fig. 4B). The number of NMJs in *Mfn2^H361Y/+^* muscle was not altered (Fig. 4D), however there was a clear reduction in endplate volume. This reduced endplate volume was completely suppressed by SARM1 loss (Fig. 4F), suggesting that degeneration of muscle fiber segments underlying the synapse is due to muscle fiber denervation (55). Furthermore, ultrastructural analysis of the NMJ revealed severe disorganization with loss of synaptic clefts between pre- and post-synaptic membranes in *Mfn2^H361Y/+^* rats and increased collagen fibers in the pre-synaptic space, consistent with a loss of normal synaptic innervation. In contrast, there were no ultrastructural abnormalities detected in *Sarm1^-/-^Mfn2^H361Y/+^* synapses (Fig. 4 G, H). In summary, the *Mfn2^H361Y^* mutation causes NMJ atrophy in a SARM1-dependent manner, suggesting that mitochondrial dysfunction largely leads to NMJ and axonal damage via SARM1 activation.

### Mitochondrial defects in *Mfn2^H361Y/+^* distal nerves are mitigated by *Sarm1* deficiency

MFN2 is localized to the mitochondrial outer membrane and is a key player in mitochondrial oxidative function, mitochondrial fusion and transport, ER-mitochondrial tethering, and other mitochondrial functions (3). As such, mitochondrial pathology is expected to be associated with *MFN2* mutations, and indeed the mitochondria in CMT2 patient nerves display altered morphology and distribution (2). Mitochondria are also actively recruited to synapses to support synaptic activity by maintaining local ATP synthesis (56). Therefore, to examine the effects of *Mfn2* mutation on synaptic mitochondria, we analyzed the number and morphology of mitochondria around the NMJs of lumbrical muscles in *Mfn2^H361Y/+^* rats by electron microscopy. We found a significant reduction in the number of mitochondria in axon termini of *Mfn2^H361Y/+^* rats (Fig. 5E). Furthermore, most of the pre-synaptic mitochondria that were present at these distal synapses displayed a swollen, rounded shape with either low cristae density or a complete lack of discernable cristae (Fig. 5A). Despite the intrinsic mitochondria defect in these rats, the mitochondria in *Sarm1^-/-^Mfn2^H361Y/+^* were present in normal numbers and had largely typical morphology. They were slightly swollen compared to WT (Fig. 5A-C), but cristae density was normal (Fig. 5D). We also examined mitochondria in the post-synaptic muscle and observed no abnormalities in the *Mfn2^H361Y/+^* animals (Fig. 5F, G), consistent with observations in CMT2 patient samples (2). That SARM1 loss suppresses mitochondrial abnormalities caused by a defective intrinsic mitochondrial protein, MFN2, is surprising and suggests a feedback loop between mitochondrial dysfunction and SARM1 activation triggering a cascade of increasing mitochondrial damage.

**Figure 5.**
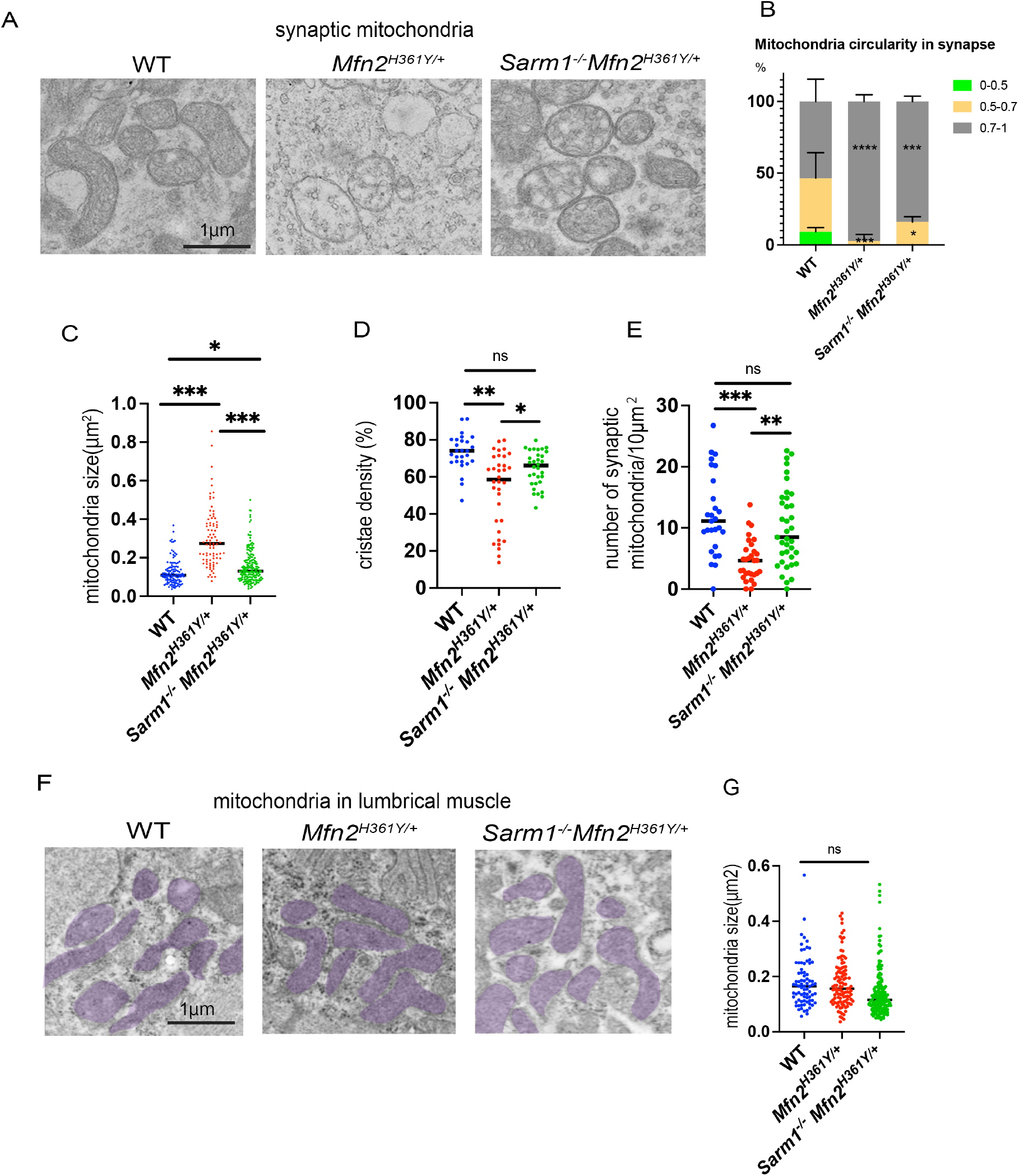
*Sarm1* deletion rescues mitochondrial defects in the synapses of *Mfn2^H361Y/+^* rats. (A) Representative images of mitochondria in synapses of WT, *Mfn2^H361Y/+^*, and *Sarm1^-/-^Mfn2^H361Y/+^* rats. (B) Percentage of elongated (circularity = 0.7-1), oval (0.5-0.7), and rounded (0-0.5) mitochondria in synapses (n =3). *p<0.05, ***p<0.005, ****p<0.001. (C) Quantification of mitochondria size in synapses (n=3). *p<0.05, ***p<0.005. (D) Quantification of cristae density of mitochondria in synapses (n =3). *p<0.05, **p<0.01. (E) Quantification of mitochondria density in synapses (n =3). **p<0.01, ***p<0.005 (F) Representative images of mitochondria in muscle from WT, *Mfn2^H361Y/+^*, and *Sarm1^-/-^Mfn2^H361Y/+^* rats. (G) Quantification of mitochondrial size in muscle (n =3).

Electron microscopic analysis of *Mfn2^H361Y/+^* sciatic and tibial nerve axons also revealed abnormal mitochondria with the same swollen, rounded shape reminiscent of human CMT2A pathology (Fig. 6A) (2). Here again a striking distal preference is exhibited in the *Mfn2^H361Y/+^* rat, as the most severe mitochondrial morphological abnormalities were observed at synapses (Fig. 5B), followed by distal and then proximal nerve segments (Fig. 6B-D). In contrast, mitochondria in *Mfn2^H361Y/+^* neuronal cell bodies appeared normal (Fig. 6E). Thus, the preponderance of mitochondrial deficits occurs in the more distal regions of the nerve including distal axons and, ultimately, at the synapse, consistent with the more severe degeneration of these structures observed in CMT2A neuropathy. The pathological mitochondrial abnormalities were again mitigated by loss of *Sarm1*, with the mitochondrial cristae density in *Sarm1^-/-^Mfn2^H361Y/+^* animals largely normal (Fig. 6D). However, the majority of *Sarm1^-/-^Mfn2^H361Y/+^* mitochondria still exhibited an aberrantly rounded shape in distal nerves (Fig. 6C).

**Figure 6.**
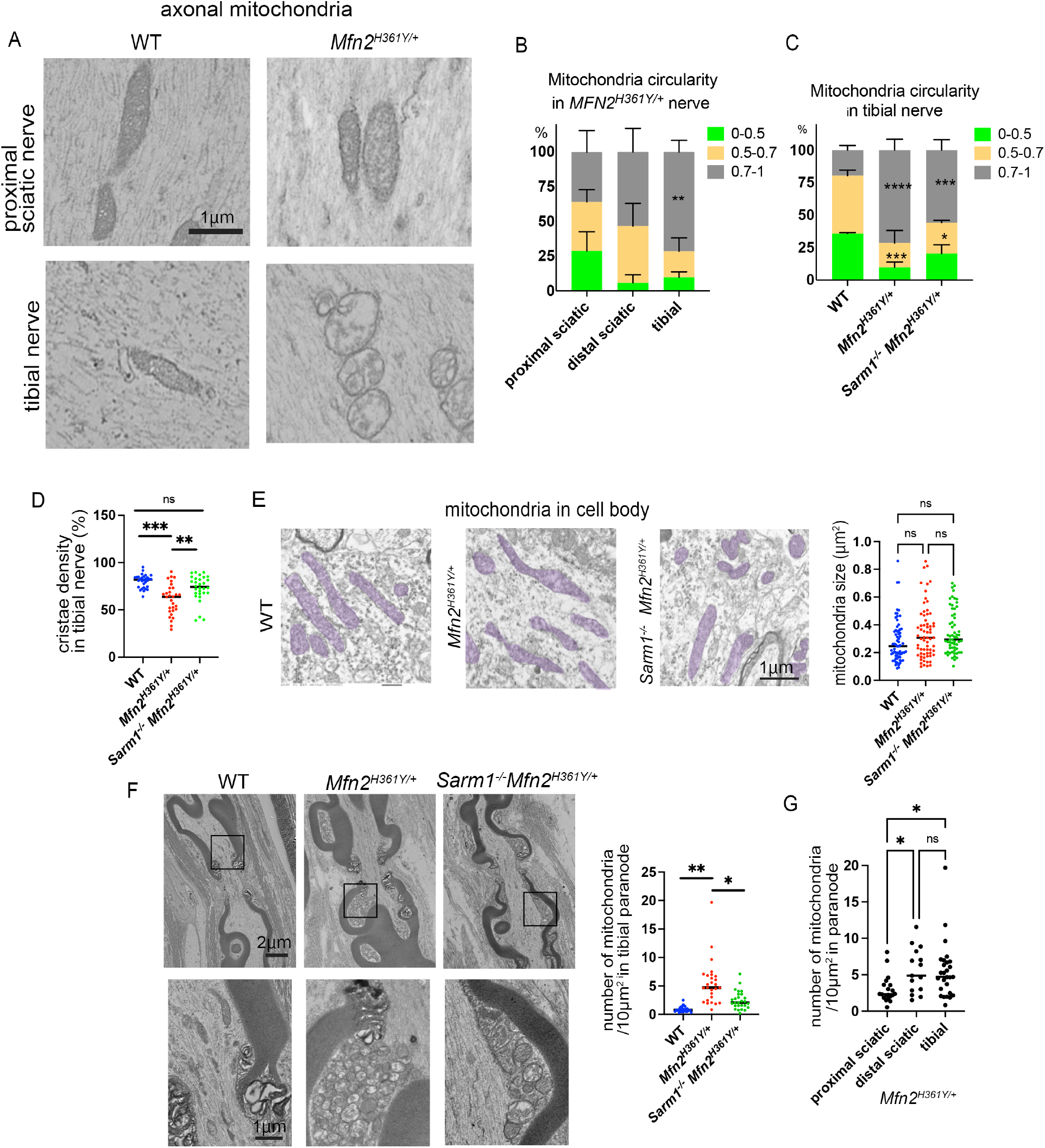
*Sarm1* deletion rescues mitochondrial defects in the axons of *Mfn2^H361Y/+^* rats. (A) Electron microscope images of proximal sciatic and tibial nerves from WT and *Mfn2^H361Y/+^* rats showing representative axonal mitochondria. (B) Percentage of elongated, oval, and rounded mitochondria categorized by their circularity in three different *Mfn2^H361Y/+^* nerves (n=3). **p<0.01, comparison between proximal sciatic and tibial. (C) Percentage of elongated, oval, and rounded mitochondria categorized by their circularity in WT, *Mfn2^H361Y/+^*, and *Sarm1^-/-^Mfn2^H361Y/+^* tibial nerve axons (n =3-4). *p<0.05, ***p<0.005, ****p<0.001, comparison between WT and mutant. *Mfn2^H361Y/+^* data is repeated from B for comparison. (D) Cristae density of mitochondria in WT, *Mfn2^H361Y/+^*, and *Sarm1^-/-^Mfn2^H361Y/+^* tibial nerve axons (n =3-4). **p<0.01, ***p<0.001. (E) Electron microscope images of mitochondria in WT, *Mfn2^H361Y/+^*, and *Sarm1^-/-^Mfn2^H361Y/+^* motor neuron cell bodies (n=3). (F) Representative electron microscope images of longitudinal axons in tibial nerves showing nodes of Ranvier. Lower images are magnified views of frames in upper images showing aggregated mitochondria. Graph represents number of mitochondria per 10μm^2^ of paranode in WT, *Mfn2^H361Y/+^*, and *Sarm1^-/-^Mfn2^H361Y/+^* axons (n =3). *p<0.05, **p<0.01 (G) Number of mitochondria per 10μm^2^ area of paranode of proximal sciatic nerve, distal sciatic nerve, and tibial nerve in *Mfn2^H361Y/+^* rats (n =3). *p<0.05.

Mitochondria play a central role in energy generation, metabolite synthesis, and calcium buffering (57). Thus, their proper localization in neurons is essential for normal function, and their sparsity in distal regions of the nerve and at the NMJ is likely involved in the pathophysiology of CMT2A. The scarcity of mitochondria in these regions led us to examine their distribution in axons from sciatic and tibial nerves. We found large, abnormal accumulations of mitochondria around the juxta paranode in *Mfn2^H361Y/+^* axons (Fig. 6F), particularly in the distal regions (Fig. 6G). Almost all of the mitochondria in these abnormal accumulations exhibit an abnormally rounded shape (Fig. 6F). Consistent with the above results, the loss of SARM1 decreased the numbers of mitochondria that accumulated in the juxta paranodal regions of *Sarm1^-/-^Mfn2^H361Y/+^* axons (Fig. 6F). The suppression of these mitochondrial abnormalities by loss of SARM1 again suggests an interplay between mitochondrial health and SARM1 axonal energetic regulation.

### Mitochondrial transport defect in *Mfn2^H361Y/+^* axons prevented by *Sarm1* knockout

MFN2 is necessary for mitochondrial transport in cultured embryonic sensory neuron axons (58), and mitochondrial density is decreased at more distal sites in the *Mfn2^H361Y/+^* rat, suggesting mitochondrial transport may be affected in these animals. To investigate the effect of the *Mfn2^H361Y^* mutation on mitochondria motility, we infected mito-GFP lentivirus into cultured adult rat DRG neurons to visualize mitochondria in axons. Time-lapse imaging and kymograph analysis demonstrated a significant decrease in mitochondrial motility in *Mfn2^H361Y/+^* axons (Fig. 7A C), with a significant increase in the percentage of stationary mitochondria and a decreased velocity of the motile mitochondria (Fig. 7A, 7B). By contrast, the number of stationary mitochondrial and mitochondrial motility in *Sarm1^-/-^Mfn2^H361Y/+^* axons were similar to that observed in WT neurons (Fig. 7B, 7C), indicating that SARM1 activity influences axonal transport and/or the ability of mitochondria to engage the transport machinery. Taken together, these results suggest that SARM1 activity is primarily responsible for the mitochondrial motility defects in axons from the *Mfn2^H361Y^* mutant.

**Figure 7.**
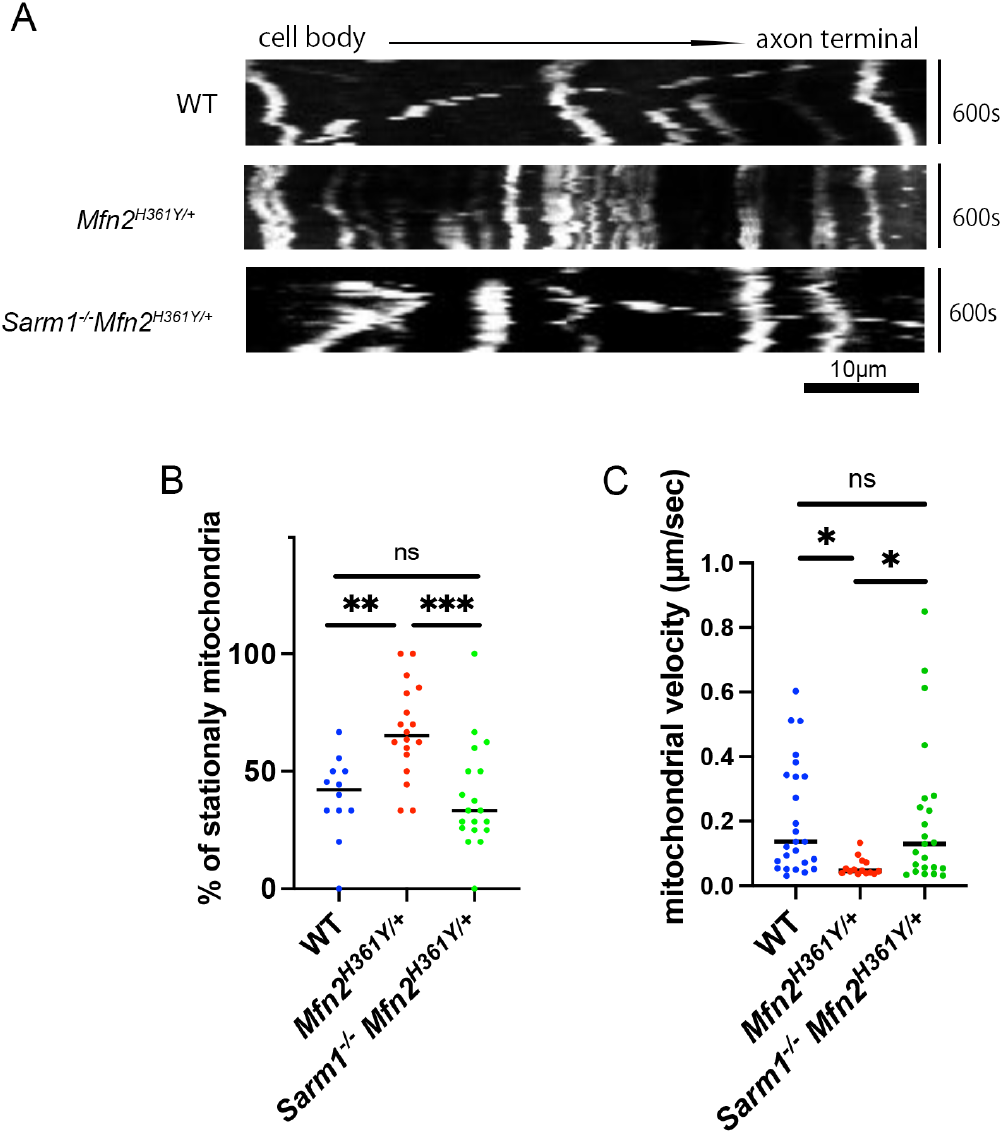
*Sarm1* deletion rescues mitochondria motility in axons of *Mfn2^H361Y/+^* rats. (A) Representative kymographs of mitochondrial movement in DRG axons. (B) Percentage of stationary mitochondria in WT, *Mfn2^H361Y/+^*, and *Sarm1 Mfn2^H361Y/+^* DRG axons (n = 3). **p< 0.01, ***p< 0.005. (C) Motile mitochondrial velocity in WT, *Mfn2^H361Y/+^*, and *Sarm1^-/-^Mfn2^H361Y/+^* DRG axons(n=3). *p<0.05.

## Discussion

Charcot-Marie-Tooth (CMT) type 2A is a debilitating axonal neuropathy with prominent axon loss and muscle wasting that is caused by mutations in the mitochondrial fusion protein *MFN2*. Here we characterize a recently described rat CMT2A model (35) and demonstrate that it displays both the hallmark neuropathological features and prominent mitochondrial abnormalities observed in the human disease. SARM1, the central executioner of pathological axon degeneration, is an inducible NAD^+^ hydrolase that can be activated by mitochondrial dysfunction, and we therefore hypothesized that SARM1 mediates pathology in CMT2A. We generated *Sarm1* knockout rats as well as *Sarm1, Mfn2* double mutant rats and find that deletion of *Sarm1* rescues the axonal, synaptic, and muscle defects in this CMT2A model. Hence, SARM1 inhibition is an exciting therapeutic candidate for CMT2A. Surprisingly, not only did *Sarm1* KO rescue neuropathological phenotypes, but it also suppressed many of the morphological characteristics of mitochondria associated with mutation of *MFN2*. These findings identify a positive feedback loop in which dysfunctional mitochondria activate SARM1 and activated SARM1 exacerbates mitochondrial dysfunction. This discovery has important implications for the potential efficacy of SARM1 inhibition in the many neurodegenerative diseases with prominent mitochondrial dysfunction. SARM1 inhibition may not only block downstream axon loss, but may also mitigate upstream mitochondrial pathology for which no current treatments exist.

### SARM1 is an essential driver of neuropathology in CMT2A

CMT2A is a debilitating neuropathy that usually leaves patients non-ambulatory as children. It is the most common of the axonal CMT neuropathies, and is caused by mutations in the mitochondrial fusion protein *MFN2*. As such, the proximate molecular cause is well-understood—loss of MFN2 function disrupts mitochondrial fusion as well as many mitochondrial functions including mitophagy, transport, and inter-organelle communication. The fundamental pathological defect in CMT2A is a distal-predominant dying back motor and sensory axonopathy with marked muscle atrophy and associated weakness. How do these mitochondrial defects result in dying back axon loss? Two prominent explanations are that dysfunctional mitochondria either cannot meet the axon’s high energetic demands and/or that they fail to sufficiently buffer calcium (3, 57). These are both reasonable possibilities since loss of ATP and calcium influx are two important drivers of axon loss. According to these explanations, the distal predominance of the pathology is attributed to defects in mitochondrial transport. However, the identification of SARM1 as the central executioner of axon degeneration in response to pathological insults including mitochondrial damage raised an alternate hypothesis—that dysfunctional mitochondria in CMT2A activate SARM1, and that SARM1 activity, rather than mitochondrial-autonomous defects, triggers the progressive, dying-back axonopathy. Here we provide strong support for this latter hypothesis.

Our analysis of pathology in the recently described *Mfn2^H361Y^* knock-in rat model of CMT2A (35) demonstrated good concordance with the human disease, i.e. progressive distal-predominant axonopathy with muscle atrophy and NMJ defects. Prominent staining for the axon regeneration marker STMN2 (41) also provided strong evidence for compensatory axon regeneration in older nerves. To test the contribution of SARM1 activity to these phenotypes, we used CRISPR-mediated gene editing to generate a *Sarm1* knockout rat and then produced *Mfn2^H361Y^,Sarm1* double mutant rats. The absence of SARM1 dramatically suppressed the pathological defects associated with *Mfn2* mutation, rescuing the hallmark axonal, synaptic, and muscle abnormalities. Hence, although compromised mitochondrial function initiates disease, SARM1 is the ultimate driver of neuropathology. If these results hold true in patients, then they have important therapeutic implications, as inhibition of SARM1 would be predicted to dramatically slow disease progression. Both small molecule and gene therapy SARM1 inhibitors block axon loss *in vitro* and *in vivo* (15, 26, 27), and AAV-mediated delivery of a SARM1 dominant negative transgene ameliorates behavioral and pathological phenotypes in another model of a human motor neuropathy (19).

How might *MFN2* mutations lead to SARM1 activation? While not yet fully understood, a strong hypothesis emerges from the mechanistic details of SARM1 activation, recently elucidated by others and ourselves. The potent SARM1 antagonist and NAD^+^ synthetase, NMNAT2, is a highly labile protein whose levels are maintained through continuous transport from neuronal cell bodies to axons (59, 60), such that NMNAT2 levels are highly sensitive to disturbances in axon integrity, transport, or energetics, especially in the most distal portions of long axons. In healthy axons, NMNAT2 suppresses SARM1 activation by locally generating NAD^+^ from its precursor NMN. SARM1 is a metabolic sensor regulated by the competitive binding of NAD^+^ and NMN to an allosteric binding site in its N-terminal autoinhibitory domain. An increase in the NMN to NAD^+^ ratio, as occurs when NMNAT2 axonal levels are low, leads to increased NMN occupancy and induces SARM1 activation (6). Pro-degenerative stimuli that activate SARM1 include different types of mitochondrial poisons (22, 23), and these toxins induce the loss of NMNAT2 (24, 25), possibly due to defects in NMNAT2 transport. However, while reduced NMNAT2 transport to distal axons may be a straightforward explanation linking mitochondrial dysfunction and SARM1 activation, we suggest a further, complementary mechanism. Since NMNAT2 requires ATP to synthesize NAD^+^ from NMN, impaired mitochondrial oxidative phosphorylation that impacts local ATP levels could also hinder NAD^+^ synthesis, thus promoting SARM1 activation even in the presence of NMNAT2. These proposed mechanisms are not specific to CMT2A and MFN2, and suggest that SARM1 activation is a common feature of many neurological disorders with mitochondrial dysfunction. If so, SARM1 inhibition may be a broadly applicable therapeutic strategy.

### A mitochondrial/SARM1 feedback loop exacerbates mitochondrial dysfunction in CMT2A

Loss of SARM1 not only suppressed pathological phenotypes in this CMT2A model, but also prevented many of the mitochondrial defects. This is an extremely surprising result, as pathogenic MFN2, a mitochondrial outer membrane protein, is still present. If mitochondrial defects were due solely to the loss of MFN2 biochemical function, then those defects should not be suppressed by loss of SARM1. Instead, our findings imply that activated SARM1 feeds back onto mitochondria and exacerbates the mitochondrial phenotypic abnormalities. It is instructive to consider which phenotypes were and were not suppressed by loss of SARM1. In *Mfn2* mutant neurons, mitochondria take on a much more rounded shape than in wild type neurons, consistent with the known roles of mitochondrial fusion as a direct regulator of mitochondrial shape (4). Although there is a modest trend toward improvement of this phenotype with loss of SARM1, the difference is not statistically significant. In contrast, *Sarm1* KO significantly suppresses the decrease in mitochondrial cristae density observed in distal nerve and synapses of *Mfn2* mutant animals. Cristae density is a morphological defect that correlates with electron-transport chain efficacy (61), and is therefore reflective of mitochondrial function. *Sarm1* KO also fully suppresses the increase in the size of mitochondria seen in the *Mfn2* mutant. Mitochondrial swelling occurs when ion homeostasis is impaired in the mitochondrial matrix, disrupting osmotic balance and leading to water influx (62, 63). *Sarm1* KO also rescues the large decrease in the number of synaptic mitochondria, which may reflect improvements in both the morphological integrity and axonal transport of mitochondria. For instance, the reduced mitochondrial mobility and velocity observed in axons of cultured adult sensory neurons from *Mfn2^H361Y^* rats are rescued in *Mfn2^H361Y^,Sarm1* double mutant neurons. Finally, an additional mitochondrial phenotype is present in nerves of *Mfn2* mutant rats—large accumulations of mitochondria in outpouchings adjacent to nodes of Ranvier. While we do not know why mitochondria collect at this location, it is a region of the axon with high energetic requirements. This phenotype is also partially suppressed by *Sarm1* KO. Of note, all of these mitochondrial defects become more prominent as one moves distally down the nerve toward the synapse, consistent with the preferential activation of SARM1 in distal axons where short-lived NMNAT2 is first depleted. Taken together, these findings suggest that while mitochondrial morphology is surely impacted by the direct loss of MFN2 function, all other mitochondrial phenotypes result primarily from the actions of SARM1.

How might activated SARM1 disrupt mitochondria? SARM1 is tethered to the outer mitochondrial membrane and activated SARM1 cleaves NAD^+^, which is essential for mitochondrial function. SARM1 cleaves cytosolic NAD^+^, and mitochondria have the machinery to synthesize their own NAD^+^ via NMNAT3. However, mitochondria also actively import NAD^+^ from the cytosol via the transporter, SLC25A51 (64, 65), suggesting that loss of cytosolic NAD^+^ could influence NAD^+^ mitochondrial pools and disrupt NAD^+^-dependent mitochondrial activity including oxidative phosphorylation and ATP generation. In addition, SARM1 activity can lead to rapid ATP loss via inhibition of glycolysis, which depends on adequate NAD^+^ stores. Indeed, the ATP used for fast axonal transport is preferentially generated by glycolysis (66), providing another mechanism for SARM1-dependent mitochondrial transport defects and decreased numbers of synaptic mitochondria in the *Mfn2* mutant rat. Finally, SARM1 has been linked to mitophagy proteins (67), suggesting it could regulate mitochondrial health in other manners beyond the cleavage of NAD^+^.

The discovery of a mitochondrial dysfunction/SARM1 activation feedback loop has profound implications for understanding the pathogenesis of not only CMT2A, but potentially other forms of CMT2. Many CMT2 disease genes have been identified, and they can be grouped into shared functions (68). Two of these shared functions are mitochondrial homeostasis, as exemplified by mutations in *MFN2* or *OPA1*, and defects in axonal transport, typified by mutations leading to aberrant dynein and kinesin function. As demonstrated here and in previous publications, mitochondrial defects can activate SARM1 (22–25), as can disruptions to axon transport because NMNAT2 transport is necessary to maintain SARM1 autoinhibition. We suggest that a vicious cycle of mitochondrial dysfunction and SARM1 activation is initiated in CMT2 subtypes associated with disrupted mitochondrial activity or axonal transport, ultimately resulting in SARM1-dependent axon loss and CMT2 pathology. If correct, this hypothesis has two important implications. First, SARM1 inhibition may by a disease-modifying therapy for a broad class of CMT2 subtypes. Second, because mitochondrial dysfunction is a feature of many neurodegenerative disorders, this destructive feedback loop may contribute to both the axonal loss and mitochondrial dysfunction in these disparate diseases, with both phenotypes amenable to SARM1-directed therapeutics.

## Materials and Methods

### Animals

All animal experiments were performed under the direction of institutional animal study guidelines at Washington University in St. Louis. As a model of CMT2A, rats which carry the *p.His361Tyr Mfn2* mutation (referred to as *Mfn2^H361Y/+^* rats) were generated using CRISPR/Cas9 gene editing technology (35). Functional testing of these *Mfn2^H361Y/+^* rats show abnormalities in gait dynamics at 8 weeks with a lengthening of the gait cycle by 16 weeks (35). To examine the contribution of SARM1 to CMT2A pathology, we generated *Sarm1* knockout rats (referred to as *Sarm1^-/-^* rats) using the SD1 strain. We used CRISPR/Cas9, mixing gRNAs with recombinant Cas9 protein, and the resulting ribonucleoprotein particle was introduced into rat embryos to generate rats with a 110 base-pair deletion in *Sarm1* exon 2 (Supplemental fig. 1A). To confirm *Sarm1* knockout, we performed a western blot using brain tissue. The SARM1 band was detected at 73kDa in wild type rats and was not present in *Sarm1^-/-^* rats (Supplemental fig. 1B).

### NMJ staining and analysis

Rats were transcardially perfused with 20 ml of 4% paraformaldehyde (PFA) solution. Dissected lumbrical muscles were post-fixed in 4% PFA overnight and then washed with PBS for 15 min thrice. Samples were permeabilized with 2% TritonX-100/PBS (PBST) for 30 min, then blocked with 4% bovine serum albumin (BSA) in 0.3% PBST for 30 mins at room temperature. Muscles were incubated with anti-SV2 (Developmental Studies Hybridoma Bank AB2315387, 1:200) and anti-2H3 (Developmental Studies Hybridoma Bank AB2314897, 1:100) for 48 hr at 4°C. After incubation with primary antibodies, muscles were incubated with FITC rabbit anti-mouse IgG (Invitrogen A21121, 1/400) overnight at 4°C. The samples were then incubated with Alexa Fluor 568 conjugated α-bungarotoxin (BTX) (Biotium 00006, 1:500) for 2 hr at room temperature. To analyze NMJ morphology, z-stack images using a confocal microscope (Leica DFC7000T) were obtained. Maximal intensity projection images were reconstructed, and post-synapse volume and terminal axon diameter were analyzed using Imaris software.

### Immunohistochemistry

Fixed tissue was processed to make 15 μm thick cryo-sections as described (69). Slides were washed with PBS, and blocked with 4% BSA dissolved in 0.3% PBST for 30 min at room temperature. They were incubated with primary antibodies against STMN2 (41, 1:500) or laminin (Sigma Aldrich L9393, 1:1000) overnight at 4°C. Slides were then washed and incubated with species appropriate secondary antibodies for 2 hr at room temperature.

### Collagen staining with PicroSirius Red

5 μm Paraffin sections were prepared as described (70), and after deparaffinization, the sections were stained with a PicroSirius Red kit (VitroVivo Biotech) using the protocol provided. Sections were imaged using a Nikon eclipse 80i light microscope and images were analyzed using ImageJ.

### Mitochondria and NMJ ultrastructural analysis

Nerves and lumbrical muscles were processed and plastic-embedded specimens were prepared as described (70). 300–400 nm sections were collected onto copper grids and then stained with uranyl acetate and lead citrate and imaged by transmission electron microscopy (JEOL1200). The detailed morphology of NMJs and mitochondria were analyzed using ImageJ software. Mitochondrial circularity was quantified using the following formula Circularity = 4π*area/perimeter^2^, as described (71). As the value approaches 0.0, it indicates an increasingly elongated polygon. A circularity value of 1.0 indicates a perfect circle.

### Nerve structural analysis by light microscopy

For light microscope analysis, plastic-embedded specimens were sectioned at 400-600 nm thickness using a Leica EM UC7 Ultramicrotome. Sections were stained with 1% toluidine blue solution. Axons were imaged using a ZEISS Axioskop, and their diameter was analyzed using a IA32 imaging analysis system. To examine the distribution of axonal diameters in nerves, at least 100 axons were measured per rat.

### Western Blotting

Brain lysates were prepared as described (9). Antibodies used: Rabbit anti-SARM1 (1:1000, 13022 Cell signaling), Mouse anti-β-actin (1:4000, A2228 Sigma Aldrich), HRP-conjugated anti-rabbit (1:5000, AB 2307391 Jackson Immuno Research), and HRP-conjugated anti-mouse (1:5000, 115-035-003 Jackson Immuno Research).

### Mass Spectrometry

Tibial nerves were homogenized in 160 μl of 50%MeOH in water, then centrifuged (15000g, 10 min). 50 μl chloroform was added into supernatant, and centrifuged again (15000g, 10 min). The clear aqueous phase was lyophilized, and metabolites were measured as described (72).

### DRG culture

96- well glass bottom tissue culture plates (Cellvis P96-0-N) were coated with poly-d-Lysine (0.1 mg/ml; Sigma) and laminin (3 μg/ml; Invitrogen) before DRG dissection. DRG were dissected from adult rats (10-12 months old) (L2-L5) and incubated with 0.5% collagenase (Gibco 17104019) at 37 °C for 1 hr. Cell suspensions were then triturated by gentle pipetting and washed 3 times with DRG growth medium (Neurobasal medium (Gibco 21103049) containing 2% B27 (Invitrogen 17504044), 100 ng/ml 2.5S NGF (Harlan Bioproducts), 1 μM 5-fluoro-2’-deoxyuridine (Sigma Aldrich 200-072-5), penicillin, and streptomycin). Cell suspensions with densities of 2.0×10^5^ cells/ml were prepared, and 1.5μl placed in the center of each well. Cells were allowed to adhere for 15 min, and then 100 μl DRG growth medium was added.

### Mitochondrial motility analysis

DRG neurons were infected with lentivirus expressing Mito-GFP (1–10 × 10^3^ pfu) at 2 days in vitro (2DIV). To analyze mitochondrial movement, Mito-GFP labeled mitochondria in distal axons were monitored using Zeiss Cell discoverer 7 at 7DIV. Frames were collected every 15 secs for 10 min. Kymographs were generated by Fiji image processing package in ImageJ and the percentage of motile mitochondria was quantified.

### Statistics

Graphpad Prism was used to perform statistical analyses by either two-tailed unpaired Student’s t-tests or one-way ANOVA with Dunnett’s multiple comparison tests. A p-value <0.05 was considered significant.

## Acknowledgments

We would like to thank members of the DiAntonio and Milbrandt labs for their thoughtful feedback on this work. We would like to thank Cassidy Menendez, Rachel McClarney, and Alicia Neiner for their technical support. We also thank the Washington University Core for Cellular Imaging (WUCCI) for their technical support, expertise, and training. We gratefully acknowledge the CMT Association for providing the CMT2A mutant rats. We also thank the Genome Engineering & Stem Cell Center (GESC@MGI) at Washington University for generating the *Sarm1* KO rats.

---

**Supplemental figure 1. Generation of *Sarm1^-/-^* rats.**

(A) The scheme indicates the deletion of *Sarm1* exon 2 in *Sarm1*^-/-^ rats. (B) Western blot analysis to detect SARM1 expression in the brains of WT and *Sarm1*^-/-^ rats. (C) Toluidine blue stained cross sections of sciatic nerve from WT and *Sarm1*^-/-^ rats. Right panels present nerves at 7days after transection, and left panels presents control nerves from the contralateral side.

