## Supplementary figures and images for "SARM1 promotes axonal, synaptic, and mitochondrial pathologies in Charcot-Marie-Tooth disease type 2A"

### supplement figure1

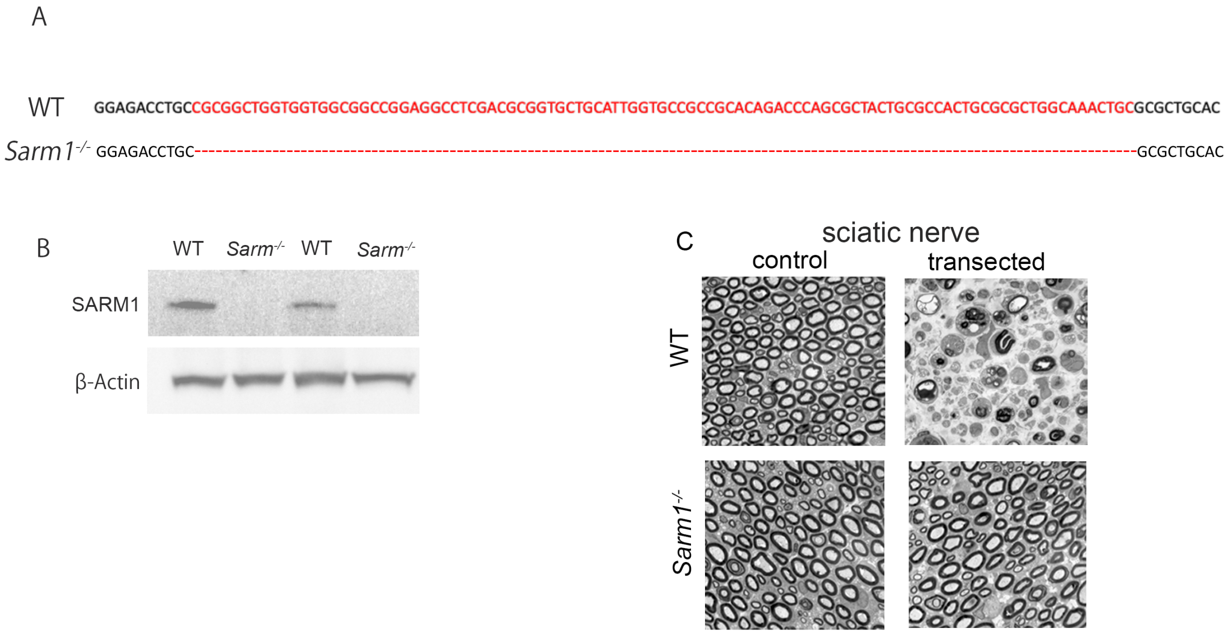
